# HYPO: A database of hypothetical human proteins

**DOI:** 10.1101/202887

**Authors:** Vijayaraghava Seshadri Sundararajan, Girik Malik, Johny Ijaq, Anuj Kumar, Partha Sarathi Das, Shidhi P.R, Achuthsankar S Nair, Prashanth Suravajhala, Pawan K Dhar

**Affiliations:** Bioclues.org, Kukatpally, Hyderabad 500072, India; Environmental Health Institute, National Environment Agency, Singapore 138667; The Battelle Center for Mathematical Medicine, The Research Institute at Nationwide Children’s Hospital, Department of Pediatrics, The Ohio state University, USA; Department of Zoology, Osmania University, Hyderabad 50007, India; Uttarakhand Council for Biotechnology (UCB), Prem Nagar, Dehradun-248007, India; Bioinformatics Infrastructure Facility, Department of Microbiology, Vidyasagar University, Midnapore 721102, West Bengal, India; Department of Computational Biology and Bioinformatics, University of Kerala, Thiruvanantapuram 695581, Kerala, India; Department of Biotechnology and Bioinformatics, Birla Institute of Scientific Research, Statue Circle, Jaipur 302001, Rajasthan, India; School of Biotechnology, Jawaharlal Nehru University, New Delhi 110067, India

## Abstract

All annotated genes were once hypothetical or uncharacterized. Keeping this as an epilogue, we have enhanced our former database of hypothetical proteins (HP) in human (HypoDB) with added annotation, application programming interfaces and descriptive features. The database hosts 1000+ manually curated records of the known ‘unknown’ regions in the human genome. The new updated version of HypoDB with functionalities (Blast, Match) is freely accessible at http://www.bioclues.org/hypo2.

## INTRODUCTION

The advent of high-throughput genomic technologies has enabled understanding the components of the genome in a better way. Today, we can distinguish sequences that are coding, non-coding and also the ones that are not the *bona fide* genes at all, *viz.* pseudogenes. Nevertheless, there are some genes whose function remains obscure as they may not have similarities to known regions in the genome. Such known ‘unknown’ genes constituting the open reading frames (ORF) that remain in the epigenome are termed as orphan genes. The proteins that are expected to be expressed from these orphan genes but having no experimental evidence of translation are termed as ‘hypothetical proteins’ (HPs) [1]. Moving from the then early stated 1.5% protein-coding sequences to the non-coding inter-and intragenic regions, all these HPs are a part of known ‘unknowns’ with undetermined role [2, 3]. Recently, the role of orphan genes and HPs has been deliberated in lieu of their role as artefacts and non-coding elements [4]. Furthermore, there are evolutionarily conserved regions (ECR) in the form of HPs which could be potential candidates for experimentation [5]. In addition, studies on structural aspects have led into determining tertiary structures of HPs based on geometrical, biophysical and biochemical studies which further emphasize the need for descriptors of these sequences [6]. Since 2006, we have been updating our primary database of HPs in human. Many of the reviews focusing on annotation from sequence/literature based searches [7], structural genomics [8] and functional genomics based approaches [9] have been well documented even as they constitute substantial part of human proteome. Due to lack of experimental confirmations regarding their molecular function, some of these variants are known as KIAA in bacteria while in eukaryotes they are tagged as 'unknown' as 'uncharacterized' with many accessions starting with prefix, “XP,” meaning predicted. Recently, we have reported making novel proteins from non-coding and less *bona fide* sequences, pseudogenes [10]. With the deluge of sequencing data pertaining to human HPs, there is a need to organize them into a database for their latent use and understand prime functions associated with various pathways and diseases. This would definitely serve as ready reference to researchers interested in finding the role of candidate HPs. What remains interesting to see is that there are many different types of regulatory sequences associated with HPs in controlling gene expression eventually falling under this list.

## METHODS

### Data Extraction and Collection

Since 2006, we have been updating our primary database of HPs in human consistently [11]. The then 7540 of them are relatively deprecated with certainty and many are likely to be a part of potential duplicates, obscure list. Annotators and curators have made a magnanimous continuous effort in predicting the function of HPs and this finally resulted in 1048 manually curated proteins that have been experimentally proven / confirmed. They were included in our earlier version (Hypo DB 12_Mar_2012) of the Hypo database. In this current work, we have enhanced our database with new functionalities and core dependencies are added to the new version of HypoDB 2.0. Set of HPs from earlier version are considered to extract the latest information on 13_Jun_2016 from UniProtKB (uniprot.org/uniprot/). This resulted in a set of **1015** final HPs that are considered in this newly updated version of HypoDB named as “Hypo DB 13_Jun_2016 (1015)”. The HPs retrieved from UniProtKB are categorized as reviewed (n=923) and non-reviewed (n=92) and are tagged as UniProtKB[Swiss-Prot] and UniProtKB[TrEMBL], respectively in our DB. They form two sub-databases in Hypo; one with reviewed and the other with non-reviewed UniProtKB peptides. The total collection of these peptides is termed as ‘UniProtKB. The idea is to allow researchers to search and use the Hypo Database according to their research interest using experimentally validated sets (from Swiss-Prot) or non-experimentally validated set (TrEMBL) or use the complete set (UniProtKB)

## DATABASE DESIGN/ STRUCTURE/ORGANIZATION

### User Interface and Functionality

At the core backend, the java construction allows views of every queried sequence mapped to several databases, viz. EMBL, PIR, HPRD and other interesting applications that include descriptors for structural databases, interaction and association databases like STRING, gene ontology and KEGG pathway reference databases apart from many sequence databases (an example is illustrated in Figure 1).

**Figure 1:**
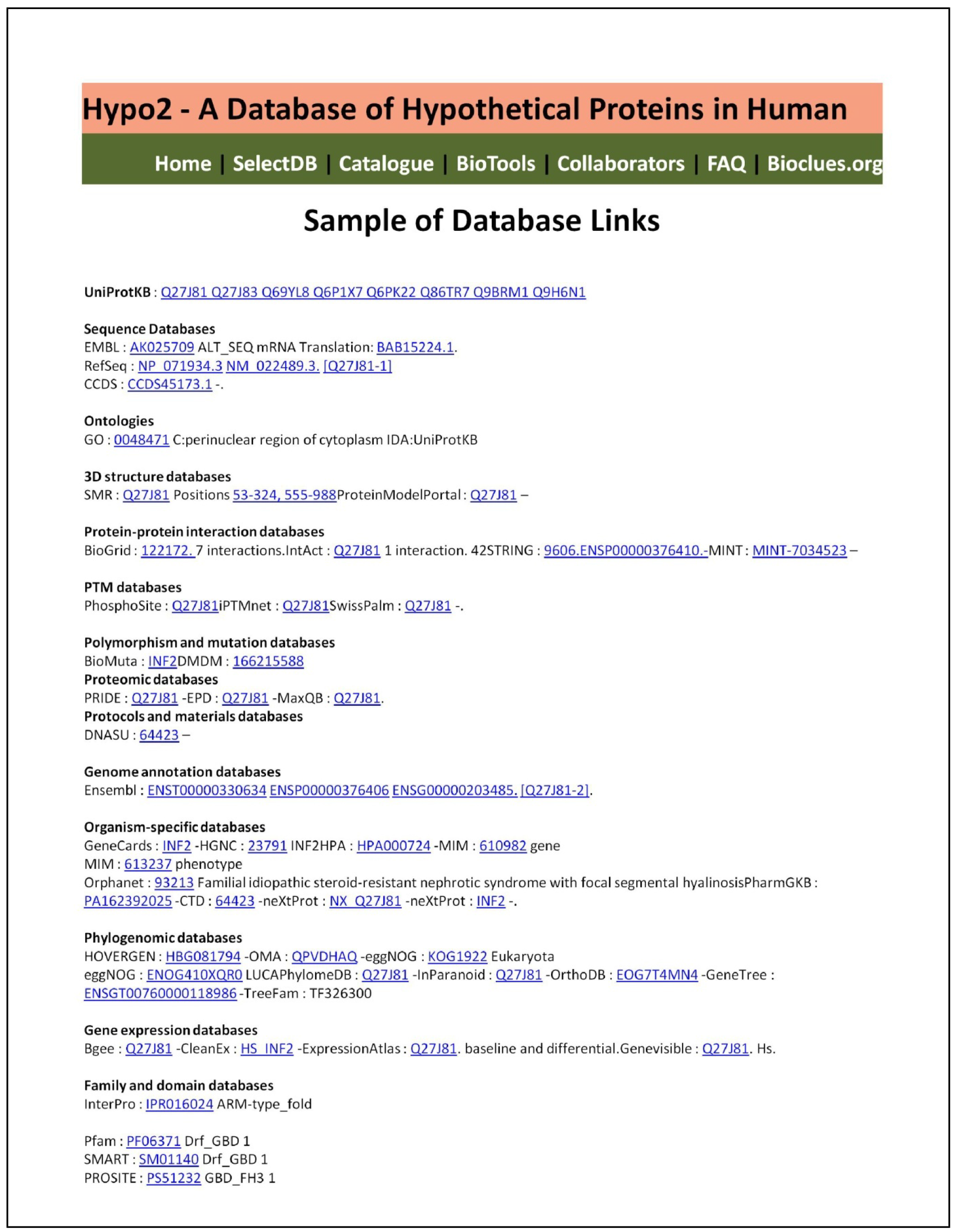
An overview of features and how the query is processed in HypoDB

The HypoDB contains primarily four application programming interfaces (API), *viz.*

(a) Quick view interface where a few modular entries can be directly retrieved from the home page of the Hypo website. HPs overlapping classes are displayed in the home page (Quick view, as shown in Figure 2). This allows user to filter the list directly according to the selected HP.
(b) Search interface (through Catalogue Sub-Menu system). Through several Catalogues, the Hypo Database is searched according to the user need and references (as detailed in a later paragraph as “Multiple search capabilities”, Figure 3).
(c) Predict sequences from BLAST I/O parser [12] (BioTool page). Users can paste or upload their sequences to be tested against the total sets of HPs (sequences) through the integrated BLAST tool.
(d) Another useful API is the core functionality with feature/annotation object model allowing the detailed view from outcome of features, while annotations can be downloaded by right click “save as” options containing ontology and featured lists. These interfaces are embedded into catalogues result page, Quick View result page, BLAST outcome pages.

**Fig.2.**
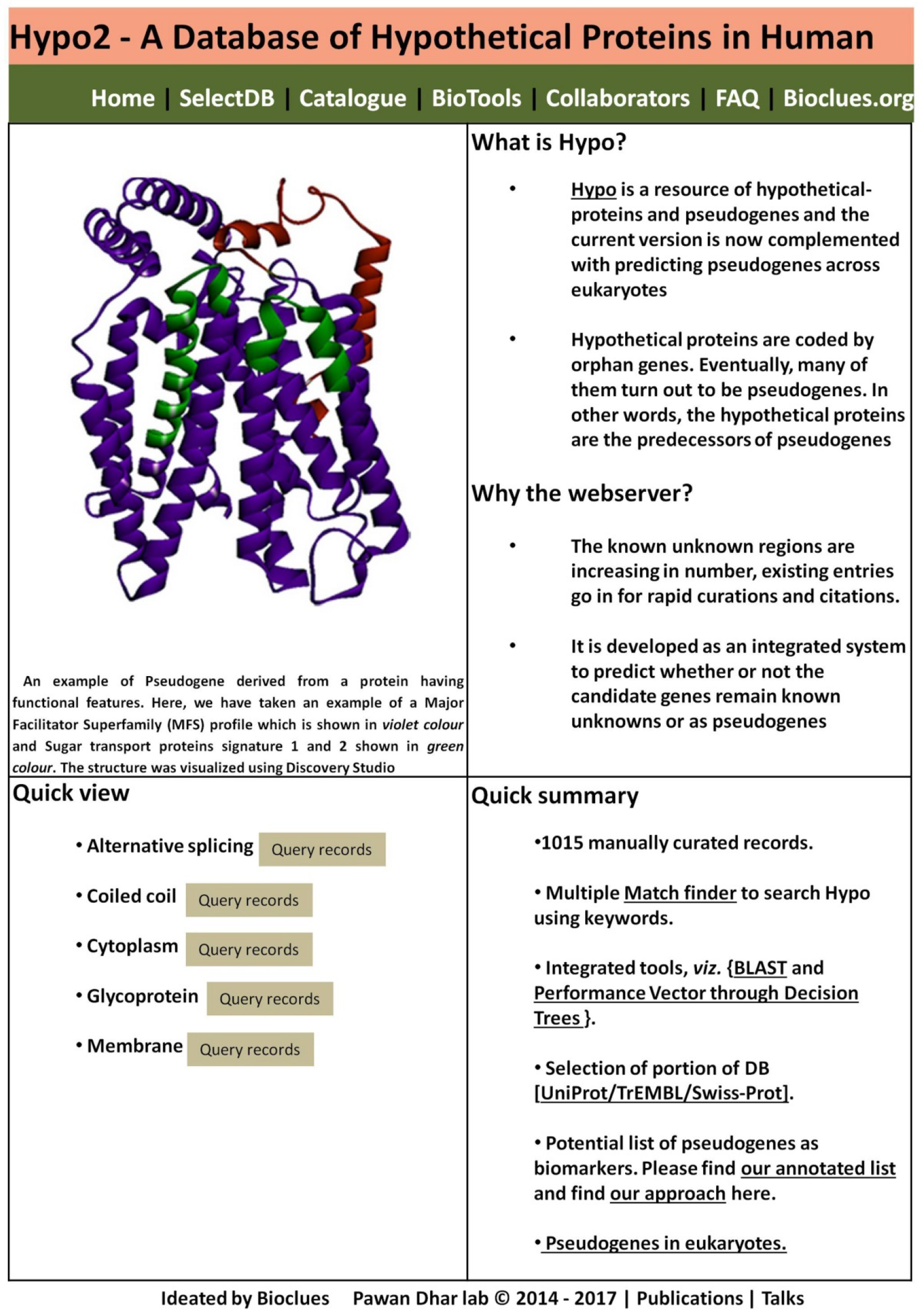
Screenshot showing the home page of the Hypo.2 database

**Figure.3.**
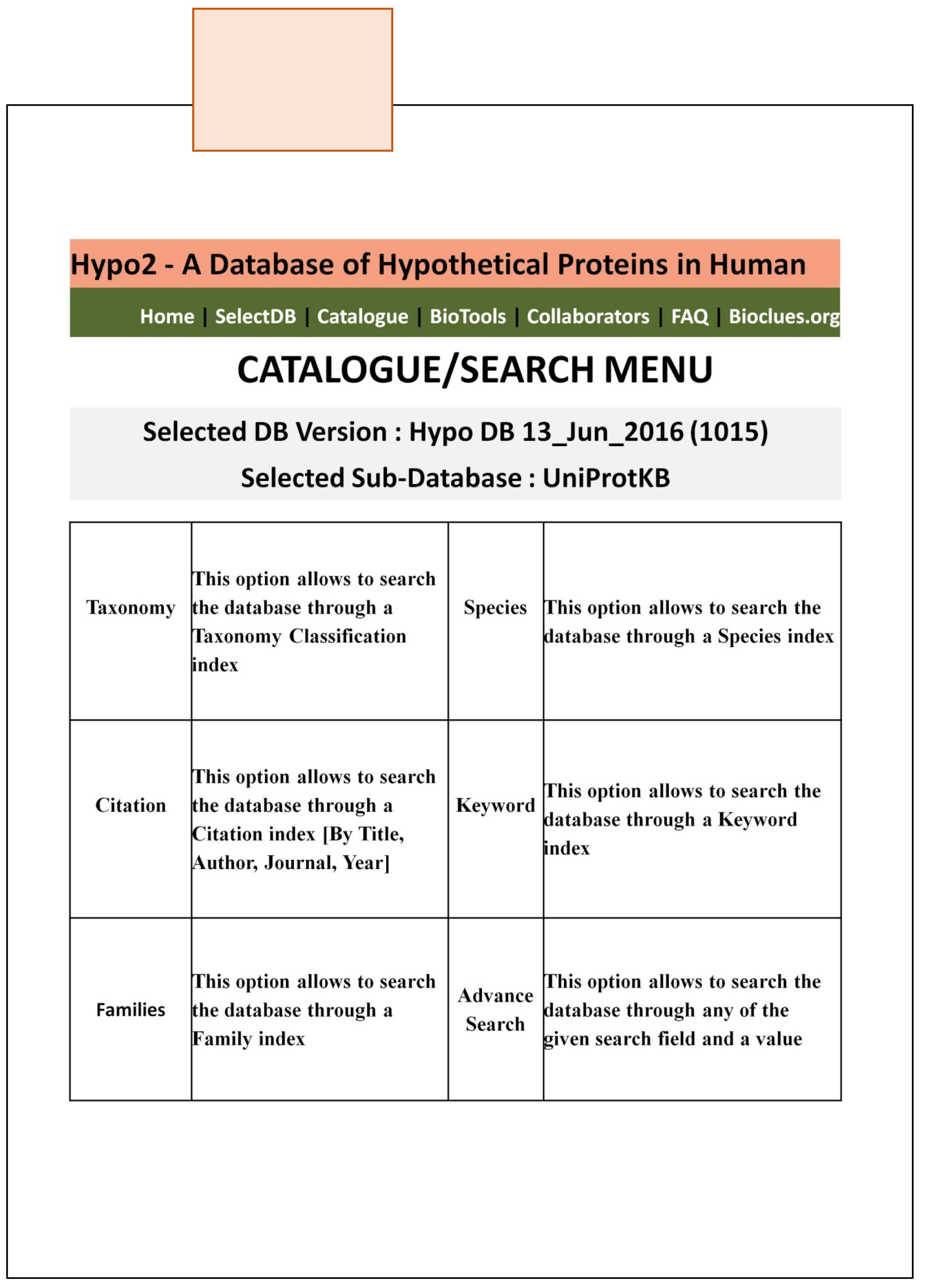
Multiple search capabilities.

### Multiple search capabilities

HYPO offers the end-user with multiple search capabilities (Figure 3). These six search tools can operate as independent search engines to interrogate the database or be executed as part of a more complex query. Five of these search utilities are based on individual catalogues that are created as vocabularies of terms from taxonomies, peptide families, species, keywords, and citations of 1015 HPs entries for easy browsing of the database.

#### Taxonomy catalogue

Organisms are classified in a hierarchical tree structure. Taxonomy database contains every node (taxon) of the tree.

#### Species catalogue

In this DB, we only considered *“Homo sapiens”* – which is set in this catalogue (the catalogue is not functional).

#### Keyword catalogue

UniProt entries are tagged with keywords that can be used to retrieve particular subsets of entries.

#### Family catalogue

In this DB, we only considered *“Homo sapiens”* – which is set in this catalogue (the catalogue is not functional).

#### Citation catalogue

UniProt maintain publications with title (RT, example: Splicing variants of BLOM7); author name (RA, examples: Abaya), journal (RL, example: Am. J. Med. Genet. A), year of publication (YR, example: 1986). In Hypo2, we have catalogued “TITLE of manuscript”, “AUTHOR Name”, “JOURNAL Name”, and “YEAR of Publication”, are separately catalogued for easy search and reference.

#### Advanced Search

The Hypo database also offers a straightforward option where user can search the database through any of the given ‘search fields’ and a ‘value’. For example, if the user selects species in ‘search field’ and *Homo sapiens* as ‘value’, it leads to a page that shows the list of all HPs described in the database. Clicking on them further shows detailed information about the selected list. This search category allows a combination of search terms, search fields and search values. Users can query the database using field names which are not listed in other catalogues. The HypoDB website also provides details on how to retrieve different components. The FAQ and help section will allow the users to get an introduction. The searches can be made in less than 10 seconds on a 2GB RAM and 2 GHz Core Duo processor.

## INTEGRATED BIO-TOOLS

**Matcher:** The Hypo database system presents a Biomarkers Matcher, along with a local gene card Matcher, which retrieves the Symbol, Description and Category of the Gene. The matcher also has an option to lookup the gene in NCBI, using the accession number. The matcher also describes the gene along with the HP and pseudogene linked to it. The result is presented in a tabular format, which can be sorted according to the needs, along with an option to perform a sub-search in the results displayed, based on any of the fields.

**BLAST** functionality is integrated in the Hypo2 system, to enable user query of any new sequence to be aligned with the Hypo database entries and get the scores for a closer match. Users also can upload their sequences in the FASTA format and blast results will be (currently) shown on the screen.

## FUTURE DEVELOPMENT

The HypoDB aims to develop APIs for implicitly searching resources linking them to other databases like NCBI Link-out. In the near future, we would like to streamline this with genomic sequences that are coding, non-coding and non-coding with coding potential, small RNAs, long non-coding RNAs that are not annotated etc. The lists of pseudogenes etc. are already under development, which will allow users to work with the concrete list. Although the Blast/GenBank parsing API is widely used, we may not use all output formats for interfacing, so a careful descriptor usage is needed even as we hope to continue the ongoing efforts with widely unsupported formats. Further, we plan to integrate analytical tools, *viz.* ClustalW, HMMER, NJplot, Hydrophobicity calculator, SignalP interfaces to the Hypo Database System. Researchers are welcome to identify niche areas and help us improve the interface. Currently the system is embedded with Java, Perl, PHP with SQL and HypoDB 3.0 version is expected by the end of 2017.

## CONCLUSIONS

The HypoDB is perhaps the only open-source HP database with a range of tools for common bioinformatics retrievals. The homepage provides access to the interfaces with all search options. We hope to ensure that this serves as a standby reference to researchers who are interested in finding candidate sequences for their potential experimental work. As we march ahead in the post genomic era, a database such as HypoDB holds importance for ascertaining factual information from annotated entries.

## ACKNOWLEDGEMENT

We thank Dr. Chanditha Hapuarachchi, Environmental Health Institute, National Environment Agency, Singapore for proof-reading the final version of the manuscript. We acknowledge Arun Gupta for his contributions in the database design (Version 1).

## Author contributions

**VSS** developed all APIs and database catalogue interfaces. **GM** provided descriptors and search interfaces. **SPR**, **AK**, **JI** and **PSD** have checked the annotations, inserted the figures and screenshots. **PS**, **AK** and **JI** wrote the preliminary version of the manuscript. **PS** and **VSS** wrote the final draft of the manuscript. **ASN**, **PKD**, and **PS** proofread the final manuscript. All authors agreed and have gone through the final version of the manuscript.

